# Chromosomal-scale *De novo* Genome Assemblies of Cynomolgus Macaque and Common Marmoset

**DOI:** 10.1101/2020.12.04.411207

**Authors:** Vasanthan Jayakumar, Osamu Nishimura, Mitsutaka Kadota, Naoki Hirose, Hiromi Sano, Yasuhiro Murakawa, Yumiko Yamamoto, Masataka Nakaya, Tomoyuki Tsukiyama, Yasunari Seita, Shinichiro Nakamura, Jun Kawai, Erika Sasaki, Masatsugu Ema, Shigehiro Kuraku, Hideya Kawaji, Yasubumi Sakakibara

## Abstract

Cynomolgus macaque (*Macaca fascicularis*) and common marmoset (*Callithrix jacchus*) have been widely used in human biomedical research. Their genomes were sequenced and assembled initially using short-read sequences, with the advent of massively parallel sequencing. However, the resulting contig sequences tended to remain fragmentary, and long-standing primate genome assemblies used the human genome as a reference for ordering and orienting the assembled fragments into chromosomes. Here we performed *de novo* genome assembly of these two species without any human genome-based bias observed in the genome assemblies released earlier. Firstly we assembled PacBio long reads, and the resultant contigs were scaffolded with Hi-C data. The scaffolded sequences obtained were further refined based on assembly results of alternate de novo assemblies and Hi-C contact maps by resolving identified inconsistencies. The final assemblies achieved N50 lengths of 149 Mb and 137 Mb for cynomolgus macaque and common marmoset, respectively, and the numbers of scaffolds longer than 10Mb are equal to their chromosome numbers. The high fidelity of our assembly is ascertained by concordance to the BAC-end read pairs observed for common marmoset, as well as a high resemblance of their karyotypic organization. Our assembly of cynomolgus macaque outperformed all the available assemblies of this species in terms of contiguity. The chromosome-scale genome assemblies produced in this study are valuable resources for non-human primate models and provide an important baseline in human biomedical research.

## Background & Summary

Cynomolgus macaque (or crab-eating macaque, *Macaca fascicularis*) and common marmoset (*Callithrix jacchus*), belonging to old world monkey and new world monkey respectively, have been widely used in human biomedical research and drug developments with expectations that they recapitulate human physiology and pathology^1^. Their genomes, consisting of 42 and 46 chromosomes in diploids^2–4^, respectively, were assembled initially using first- and second-generation sequencing technologies^5–7^. Shortread *de novo* assembly was not able to resolve complex repetitive genomic regions, and the resulting contigs tended to remain fragmentary. Techniques such as mate-pair sequencing were commonly used to join the contigs into longer scaffolds, albeit with sequence gaps in between. Long-standing non-human primate (NHP) genome assemblies released earlier used the human genome for ordering and orienting the assemblies into chromosomes, which prevents the observation of intrinsic structural differences between the primate genomes^5^. For example, a large inversion of around 20 Mb was observed in chromosome 16 of the earlier marmoset genome assembly, which should have been the result of the ‘humanization’ bias^6^.

Recent technological advancements allow us to obtain chromosome-scale assemblies without relying on existing genome assemblies, where such errors or bias can be avoided. Single-molecule long-read sequencing (Pacific Biosciences [PacBio] and Oxford Nanopore Technologies) have drastically increased the contiguity of assemblies, and chromatin contact profiling with Hi-C and other techniques such as optical mapping have paved the way to reconstructing chromosome-scale sequences. Taking advantage of these recent advancements, the genome sequences of some non-human primates (NHP) including gorilla, orangutan, and chimpanzee were largely improved^5, 7^, followed by the one for Northern white-cheeked gibbon (Bioproject accession: PRJNA369439). High-quality genome assemblies of old world monkeys were also recently reported, such as the ones for Rhesus macaque by three different research groups (PRJNA509445, PRJNA514196^8^, and PRJNA476474^9^), the ones for olive baboon (PRJNA527874^10^), golden snub-nosed monkey (PRJNA524949^11^), pygmy chimpanzee (PRJNA526933) and Francois’s langur (PRJNA488530^12^). We had also previously produced pseudochromosome assembly by using PacBio long-reads for common marmoset^6^, where ‘humanized’ sequences were still used as a reference.

In this study, we focus on cynomolgus macaque and common marmoset to establish a solid baseline for human biomedical research. We performed genome sequencing and *de novo* assembly for both species by using PacBio long-reads, along with Hi-C for chromosome-scale scaffolding through optimized Hi-C data acquisition^13^. We corrected misjoins of the scaffolds through examination of the Hi-C contact maps. We further investigated misjoins through cross-checking alternate contigs made from PacBio data and corrected them. Lastly, we assessed the quality of the resultant assemblies based on the contiguity of assembled sequences (N50), completeness of conserved proteincoding genes, and consistency to BAC clone sequences. The results indicate that our genome assemblies are of chromosomal-scale contiguity, nearly complete coverage of gene space, and highly concordant to the conventional genomic resource obtained independently to our data set. Above all, for cynomolgus macaque, our genome assembly achieved the optimal quality in comparison with other resources available for this species.

## Methods

### Sample preparation, sequencing, and *de novo* assembly

A cynomolgus monkey was purchased by Shiga University of Medical Science from Shin Nippon Biomedical Laboratories, Ltd through Angkor Primates Center Inc in Kingdom of Cambodia. The identification number is CE1976F in Shiga University of Medical Science and K150090 in Shin Nippon Biomedical Laboratories, Ltd. We followed the Reporting in Vivo Experiments (ARRIVE) guidelines developed by the National Centre for the Replacement, Refinement & Reduction of Animals in Research (NC3Rs). All animal experimental procedures were approved by the Animal Care and Use Committee of Shiga University of Medical Science (approval number: 2017-10-2 (H1)). Genomic DNA (gDNA) was extracted from the kidney of the 6-year-8-month-old female cynomolgus macaque with MagAttract HMW DNA kit (48) [Qiagen 5067-5583; Cat No 67563] according to the manufacturer’s instruction. After measurement of concentration with Qubit3 - Qubit dsDNA (BR) [Thermo Fisher Scientific] and optical density (OD) with nanoDrop 2000 [Thermo Fisher Scientific], ethanol precipitation of the extracted gDNA was performed with DNA Clean & Concentrator-10 (25) [Agilent] to condense it. Then, concentration and OD of the gDNA were again measured with NanoDrop 2000, Qubit3 - Qubit dsDNA (BR): the amount, 70.6 ug (307 ng/uL x 230 uL); 260/280, 1.87; 260/230, 2.28. Also, the successful extraction of long gDNA was confirmed by electrophoresis with TapeStation ScreenTape GenomicDNA on TapeStation [Agilent]. Using PacBio Sequel II, approximately 81× sequence data of the cynomolgus macaque was sequenced with a read N50 length of 12.05 kbp. For common marmoset, the 43× PacBio RSII read dataset from our earlier sequencing effort (PRJDB8242), was used to reassemble the genome (Fig. 1A; Table 1).

**Figure 1.**
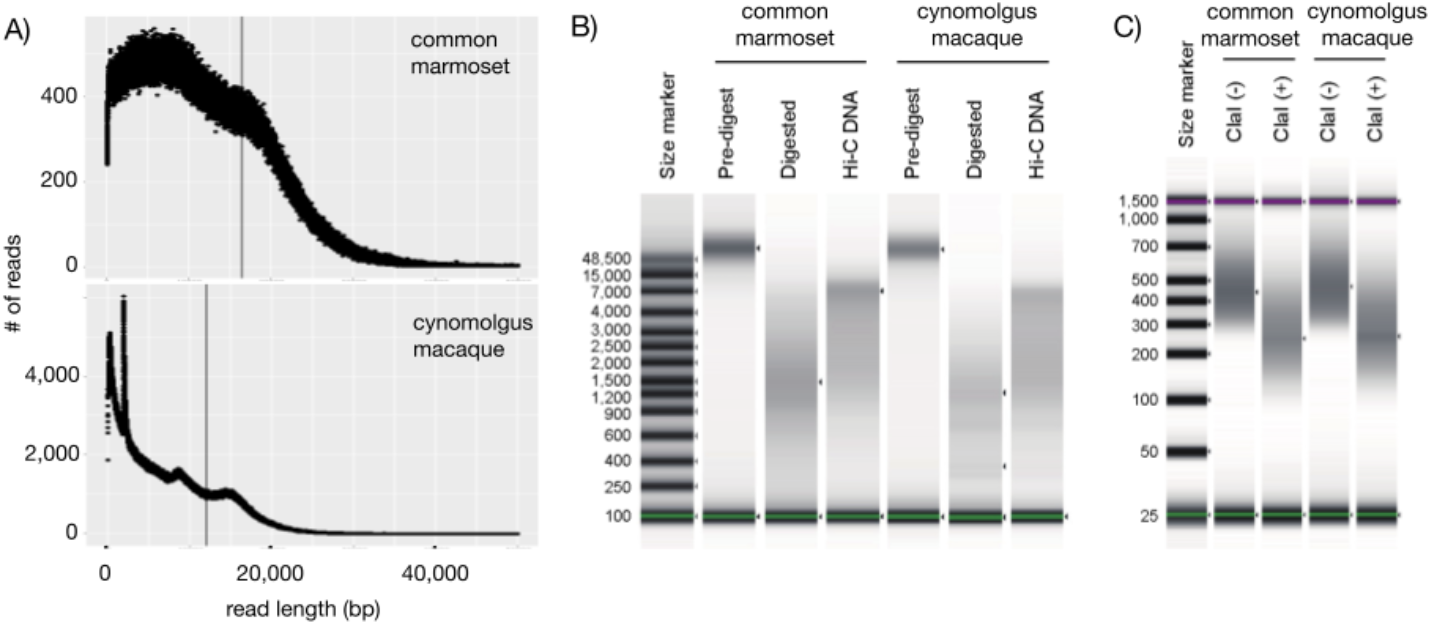
Quality assessment of the experiments. A) Read length distribution of the common marmoset and cynomolgus macaque genomes, with a vertical line representing the N50 length. B) Length distribution of the Hi-C DNA analyzed with Agilent TapeStation using the Genomic DNA ScreenTape. (C) Length distribution of the Hi-C library analyzed with Agilent TapeStation using the High Sensitivity D1000 ScreenTape.

**Table 1:** Overview of the obtained sequences

| PacBio Sequeuncing |  |  |  | iconHi-C |  | Note |
| --- | --- | --- | --- | --- | --- | --- |
| Species | Sample | Sequencer | Read coverage | Sample | Read # |  |
| Common marmoset | kidney (male) | RS II | 43x | skeletal muscle (female) | 408M reads | PacBio reads obtained from Jayakumar, V. et al 2020 |
| Crab-eating macaque | kidney (female) | Sequel II | 81x | skeletal muscle (female) | 377M reads | The samples are obtained from the same individual |

To assemble both the NHP genomes, we employed a series of assembly programs (Fig. S1). Based on the performances from the earlier marmoset genome assembly study, we chose the assemblers Flye^14^, Redbean (wtdbg2)^15^, SMARTdenovo^16^, and miniasm^17,18^. After *de novo* assembly, all the assemblies were processed by PacBio’s polishing tool, arrow, to eliminate possible sequencing errors. After considering the results from multiple *de novo* assembly tools, Redbean was chosen as the base assembler, as it produced the most contiguous assemblies for both the common marmoset (contig N50 of 8.04 Mb; Table 2) and the cynomolgus macaque (contig N50 of 5.84 Mb) genomes (Table 3).

**Table 2:**
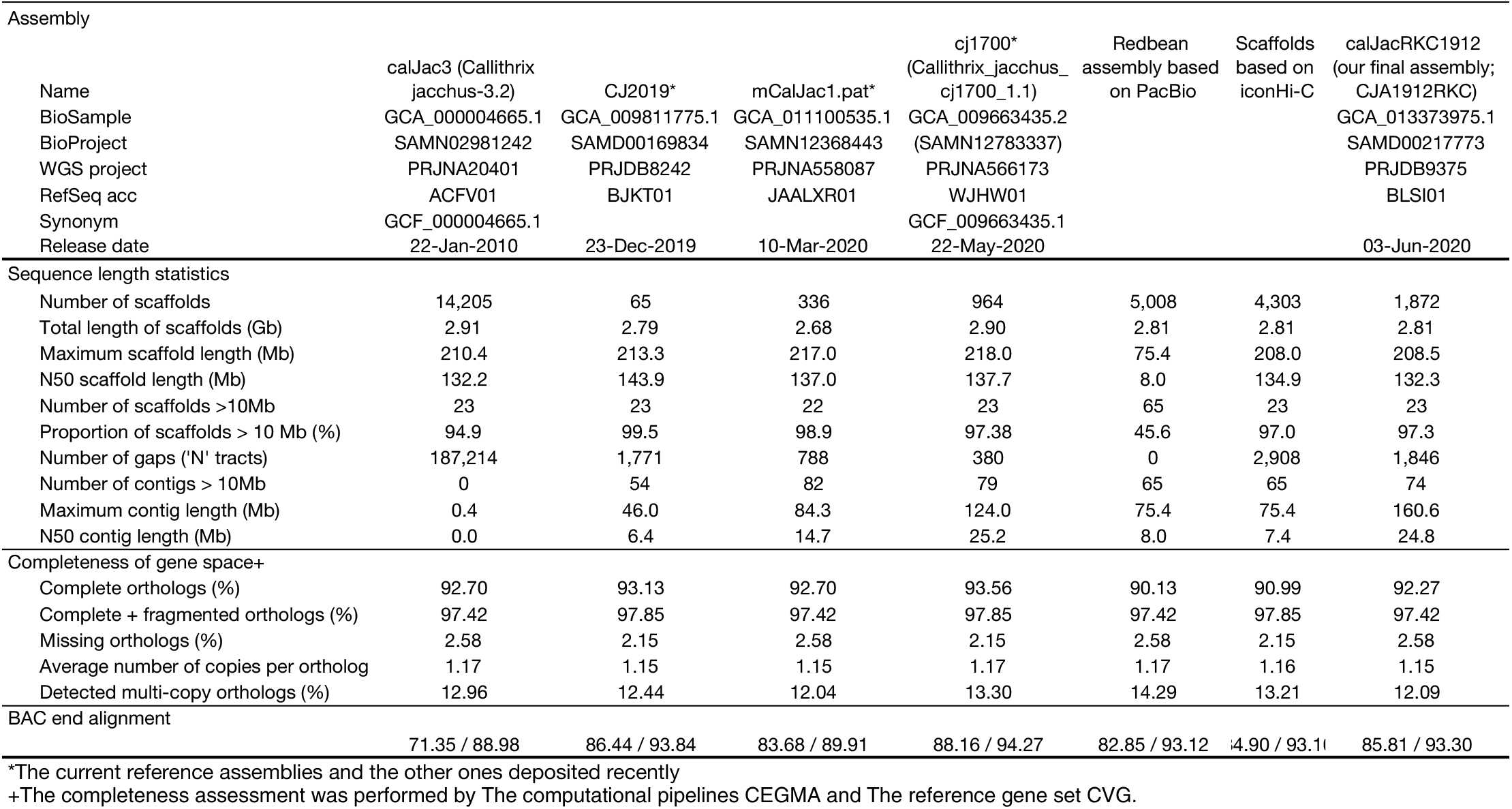
Properties of the obtained and existing assemblies of common marmoset

**Table 3:**
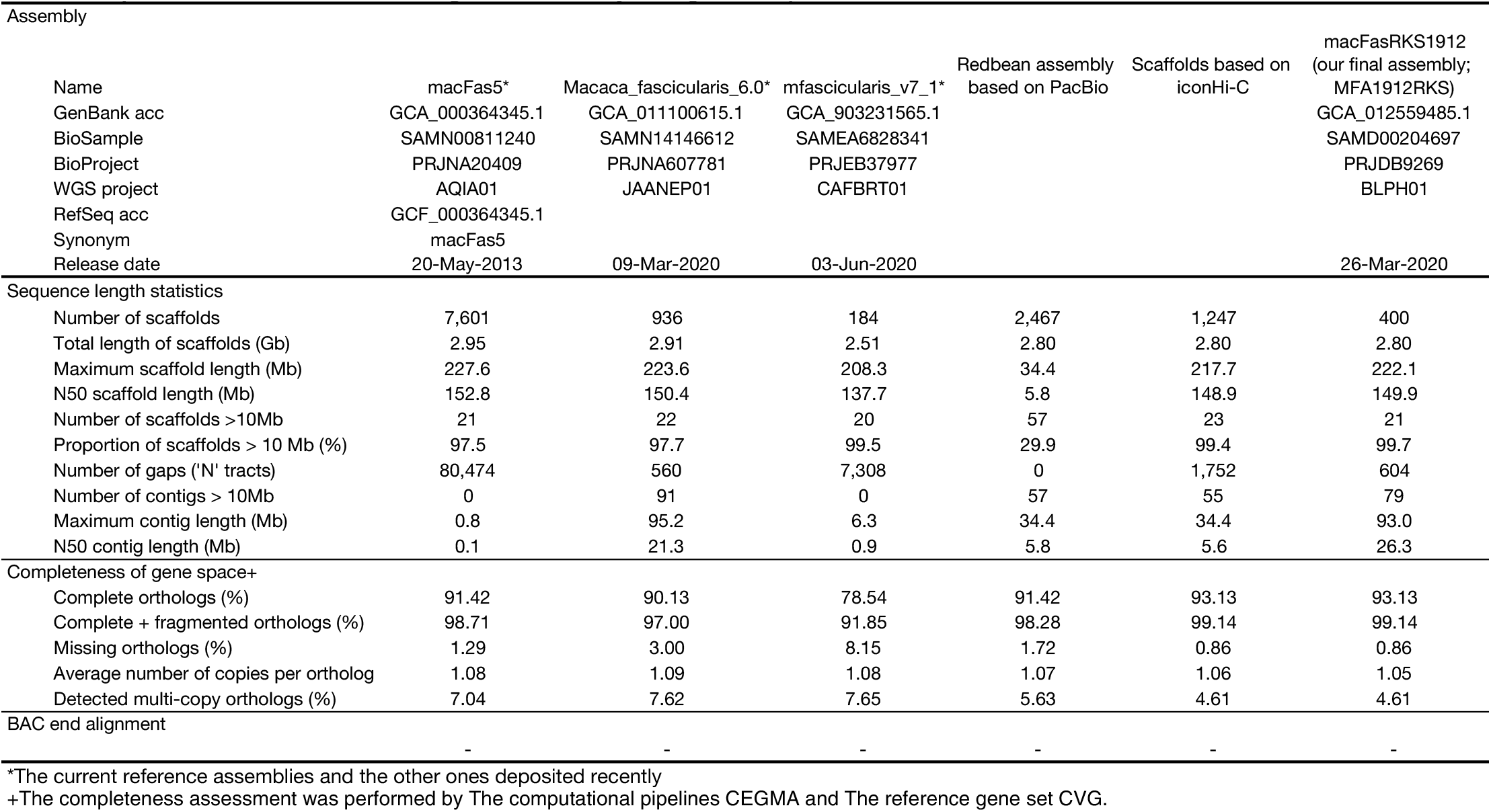
Properties of the obtained and existing assemblies of cynomolgus macaque

| Assembly |  |  |  |  |  |  |
| --- | --- | --- | --- | --- | --- | --- |
| Name | macFas5* | Macaca_fascicularis_6.0* | mfascicularis_v7_1* | Redbean assembly based on PacBio | Scaffolds based on iconHi-C | macFasRKS1912 (our final assembly; MFA1912RKS) |
| GenBank acc | GCA_000364345.1 | GCA_011100615.1 | GCA_903231565.1 |  |  | GCA_012559485.1 |
| BioSample | SAMN00811240 | SAMN14146612 | SAMEA6828341 |  |  | SAMD00204697 |
| BioProject | PRJNA20409 | PRJNA607781 | PRJEB37977 |  |  | PRJDB9269 |
| WGS project | AQIA01 | JAANEP01 | CAFBRT01 |  |  | BLPH01 |
| RefSeq acc | GCF_000364345.1 |  |  |  |  |  |
| Synonym | macFas5 |  |  |  |  |  |
| Release date | 20-May-2013 | 09-Mar-2020 | 03-Jun-2020 |  |  | 26-Mar-2020 |
| Sequence length statistics |  |  |  |  |  |  |
| Number of scaffolds | 7,601 | 936 | 184 | 2,467 | 1,247 | 400 |
| Total length of scaffolds (Gb) | 2.95 | 2.91 | 2.51 | 2.80 | 2.80 | 2.80 |
| Maximum scaffold length (Mb) | 227.6 | 223.6 | 208.3 | 34.4 | 217.7 | 222.1 |
| N50 scaffold length (Mb) | 152.8 | 150.4 | 137.7 | 5.8 | 148.9 | 149.9 |
| Number of scaffolds >10Mb | 21 | 22 | 20 | 57 | 23 | 21 |
| Proportion of scaffolds > 10 Mb (%) | 97.5 | 97.7 | 99.5 | 29.9 | 99.4 | 99.7 |
| Number of gaps ('N' tracts) | 80,474 | 560 | 7,308 | 0 | 1,752 | 604 |
| Number of contigs > 10Mb | 0 | 91 | 0 | 57 | 55 | 79 |
| Maximum contig length (Mb) | 0.8 | 95.2 | 6.3 | 34.4 | 34.4 | 93.0 |
| N50 contig length (Mb) | 0.1 | 21.3 | 0.9 | 5.8 | 5.6 | 26.3 |
| Completeness of gene space+ |  |  |  |  |  |  |
| Complete orthologs (%) | 91.42 | 90.13 | 78.54 | 91.42 | 93.13 | 93.13 |
| Complete + fragmented orthologs (%) | 98.71 | 97.00 | 91.85 | 98.28 | 99.14 | 99.14 |
| Missing orthologs (%) | 1.29 | 3.00 | 8.15 | 1.72 | 0.86 | 0.86 |
| Average number of copies per ortholog | 1.08 | 1.09 | 1.08 | 1.07 | 1.06 | 1.05 |
| Detected multi-copy orthologs (%) | 7.04 | 7.62 | 7.65 | 5.63 | 4.61 | 4.61 |
| BAC end alignment |  |  |  |  |  |  |
|  | - | - | - | - | - | - |

\*The current reference assemblies and the other ones deposited recently
+The completeness assessment was performed by The computational pipelines CEGMA and The reference gene set CVG.

### Hi-C data acquisition and scaffolding

Hi-C libraries were constructed with the restriction enzyme DpnII following the iconHi-C protocol^13^, using the skeletal muscle tissues of an adult female marmoset and a cynomolgus macaque that were kept frozen in −80°C after dissection and snap-freezing in liquid nitrogen. Fixed tissue materials containing 2 μg of DNA were used for the preparation of Hi-C DNA by *in situ* restriction digestion and ligation. Library preparation was performed using 1 μg of Hi-C DNA with 5 cycles of PCR for the marmoset and 6 cycles of PCR for the cynomolgus macaque. Quality control of the Hi-C DNA and the Hi-C library was performed as described in the iconHi-C protocol^13^. Quality control of the Hi-C DNA showed an expected pattern of a shift in size – shortening after digestion and elongation after ligation, indicating successful preparation of the Hi-C DNA (Fig. 1B). Quality control of the Hi-C library by restriction digestion confirmed the existence of expected ligation junction sequences inside the library molecules at a high proportion and successful generation of the library (Fig. 1C).

Sequencing to obtain Hi-C reads was performed on an Illumina HiSeq X in paired-ends with 151 cycles. The obtained reads were processed using Trim Galore to remove low-quality regions and adapter sequences. Post-sequencing quality control of the Hi-C libraries was performed as described previously^13^. Hi-C read statistics obtained by HiC-Pro^19^, using one million subsampled read pairs from the large scale sequencing data mapped to the PacBio contigs, confirmed high quality of the libraries while showing a high proportion of valid interaction read pairs, a low proportion of invalid ligation products (dangling end pairs), and a low proportion of contiguous restriction fragments (re-ligation pairs) (Table S1). Approximately 408 and 377 million Hi-C reads for the marmoset and the cynomolgus macaque were mapped to the Redbean contig sequences of the common marmoset and the cynomolgus macaque respectively using Juicer^20^. With the Juicer output files, Hi-C scaffolding was performed using 3d-dna^21^. Inversions and misjoins in the assemblies that occurred during the Hi-C scaffolding process were corrected by using Juicebox based on the frequency of Hi-C contacts^22^.

As a result of Hi-C scaffolding, the N50 lengths increased from approximately 8 to 135 Mb for the marmoset (Table 2) and from 5.8 to 149 Mb for the cynomolgus macaque (Table 3), and the number of scaffolds longer than 10 Mb presumed to be at the chromosome level became close to the actual number of chromosomes. For the cynomolgus macaque, two pairs of the scaffold sequences longer than 10 Mb were merged into two pseudo-chromosomes, referred to as chromosomes 2 and 8 later, based on alignment concordance with the 3d-dna scaffold sequences of alternate assemblies, as well as the previous reference sequence.

### Misjoin detection and gap-filling

All the alternate assemblies were also independently scaffolded using 3d-dna. These were later aligned against the Hi-C scaffolds from Redbean assembly to observe whether any misjoins were introduced by Hi-C scaffolding. Because the contiguity profiles of the assemblies are different across the employed assemblers, contigs which are broken into two or more sequences in one assembly could have been assembled into a single contiguous piece in another assembler (Supplementary Fig. S1). This contiguity information from alternate assemblies was used to identify misjoins in the scaffold sequences from the Redbean assembly (Fig. 2B). When the alignments of the alternate assemblies against Redbean scaffolds were visualized using D-genies dot plots^23^, occasionally the scaffolds did not exhibit consistency in chromosomal structure among the 3d-dna scaffolds. These indicated the presence of misjoins introduced by Hi-C scaffolding (Supplementary Fig. S2).

**Figure 2.**
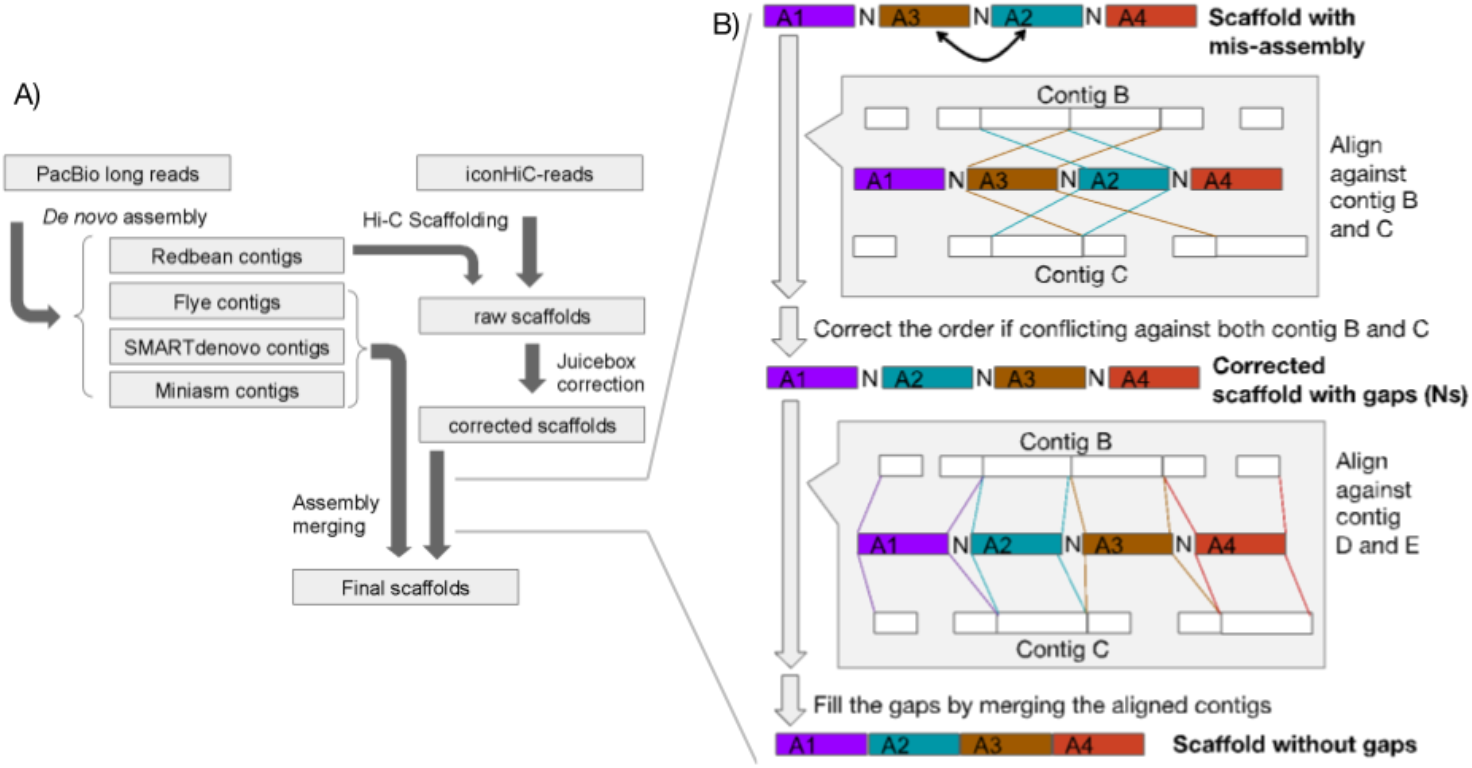
Computational steps of genome assembly. A) *De novo* assembly and Hi-C scaffolding workflow. B) Misjoin correction and gap-filling using contigs from genome assemblies using alternate assembly tools.

**Figure 3.**
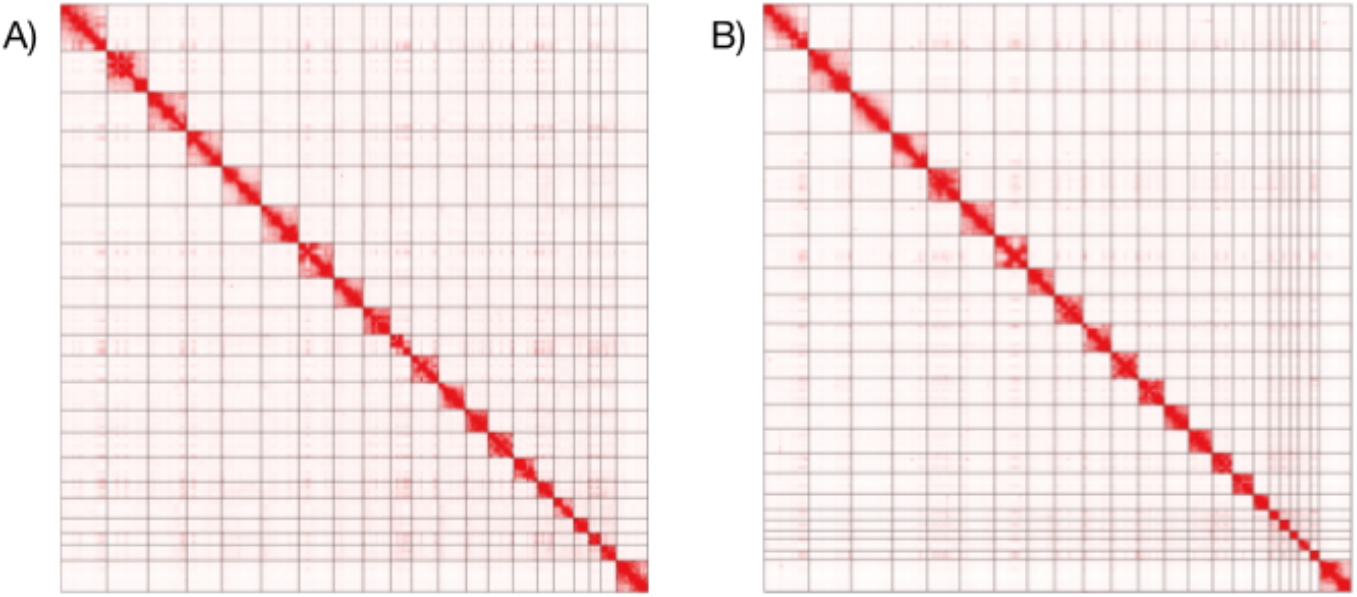
Hi-C contact map on the final assemblies. Hi-C contact maps of the pseudo-chromosome assemblies of A) common marmoset and B) cynomolgus macaque.

In case that such discrepancies in contig orders were observed in the alignments, a two-step check was performed, A) whether gaps are present in the Redbean scaffolds, and B) whether at least two of the alternate assemblies are consistent in those regions and discrepant against the Redbean scaffolds. If the above case is true, instead of re-ordering the misjoined scaffolds, we replaced the alignment block with one of the alternate assemblies making it a hybrid assembly. A similar procedure was also performed for gapfilling in regions without misjoins also. When a gap-containing region in Redbean scaffolds aligned to a gap-free region of contigs from alternate assemblies, those gap-containing regions were replaced with the corresponding blocks from alternate assemblies (Supplementary Fig. 2B).

### Data records

The N50 length of the contigs increased from 8.04 Mb to 24.82 Mb, and from 5.84 Mb to 26.27 Mb for the final assemblies of common marmoset and cynomolgus macaque genomes, respectively. The final genome assemblies displayed chromosome-sized sequences, designated here as ‘pseudo-chromosomes’, which exhibited scaffold N50 lengths of 132.27 Mb and 149.88 Mb, for the cynomolgus macaque and the common marmoset genomes, respectively (Table 2; Table 3).

The raw genome sequence data of the cynomolgus macaque are deposited at the DDBJ under the accession, DRA009584. The Hi-C raw sequence data are deposited under the accessions, DRA009641, and DRA009987 for the cynomolgus macaque and common marmoset genomes, respectively. The assembled genome sequences are deposited under the accessions, BLPH01000001-BLPH01000400, and BLSI01000001-BLSI01001872, for the cynomolgus macaque and the common marmoset genomes, respectively.

### Technical validation

#### Gene space completeness assessment

The web server gVolante^24^, in which two established pipelines CEGMA^25^ and BUSCO^26^ are implemented, was used to assess the sequence length distributions and gene space completeness in a uniform environment. The latter was based on the coverage of one-to-one reference orthologues with the ortholog search pipeline CEGMA and the gene set CVG that is specifically optimized to assess vertebrate genome sequences^27^. The analysis of gene space completeness revealed a smaller number of missing CVG genes for both species, in comparison with the scores for the genome assemblies released earlier (Table 2; Table 3).

#### BAC-end alignment

BAC-end read pairs of the common marmoset from our earlier sequencing effort^4^, were aligned against the constructed assembly using bowtie2^28^. After the misjoin detection and gap-filling procedure, the number of concordantly aligned read pairs increased by a count of 684, while the number of discordantly aligned read pairs decreased by a count of 401 (Table 2).

#### Comparison of common marmoset assembly against previous assemblies

For the common marmoset, NCBI hosts nine genome assemblies (https://www.ncbi.nlm.nih.gov/assembly/organism/9483/latest/), with cj1700 being the representative reference genome. The assembled genome in this study has a contig N50 length of 24.82 Mb in comparison to the contig N50 length of 6.38 Mb from our earlier effort (CJ2019). Our newly assembled contigs have slightly lesser contiguity than the recently submitted assembly cj1700 which has a contig N50 length of 26.62 Mb (Fig. 4A). Notably, more than half of the sequence gaps in the pseudo-chromosomal sequences of the cj1700 assembly were filled in by our assembly, implying that these assemblies are similarly useful and, in some difficult regions, complementary to each other (Table S2).

**Figure 4.**
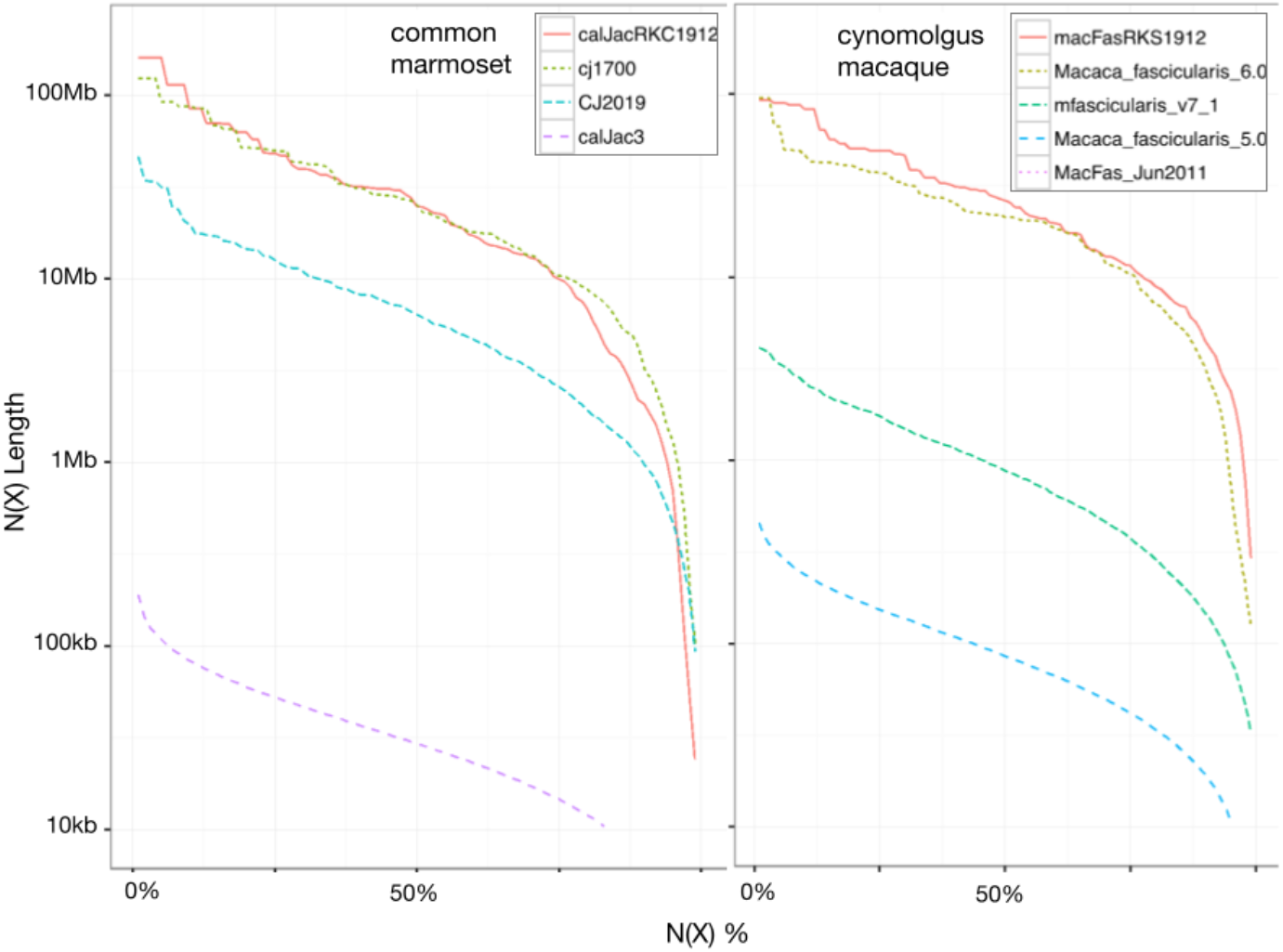
Contiguity plots of the existing and obtained assemblies. N(X) plots comparing the contiguity profiles of existing and obtained genome assemblies of A) common marmoset and B) cynomolgus macaque.

#### Comparison of cynomolgus macaque assembly against previous assemblies

NCBI hosts five cynomolgus macaque genome assemblies (https://www.ncbi.nlm.nih.gov/assembly/organism/9541/latest/), with the macFas5 genome assembly being the representative reference genome (Table 3). Three of the assemblies including the representative genome have contig N50 lengths shorter than 100 kbp. In contrast, the recently submitted Macaca_Fascicularis_6.0 assembly had produced a much larger contig N50 length of 21.34 Mb. The assembly from our present study, with a contig N50 length of 26.27 Mb, outperformed all the assemblies in terms of contiguity (Fig. 4B). Notably the coverage of orthologous genes in our assembly achieved the highest ratio, 93%, whereas the one in Macaca_Fascicularis_6.0 was 90% (Table 3). It indicates that the assembly produced here outperformed the rest not only in terms of contiguity, but also in base accuracy.

#### Comparison against other non-human primate genome assemblies

The contiguity of the two genomes assembled in this study was compared against other non-human primate genomes such as those of Rhesus macaque, Francois langur, chimpanzee, gorilla, orangutan, white-cheeked gibbon, golden snub-nosed monkey, pygmy chimpanzee, and olive baboon. Although there were three different sub-species genome assemblies for Rhesus macaque, we herein considered only the best assembly out of them in terms of contiguity. In comparison to the other non-primate genome assemblies, it was evident that the common marmoset and cynomolgus macaque assemblies produced the second and the third-best contiguities, with the Rhesus macaque genome assembly being the best among all in terms of contiguity (Supplementary Fig. S3).

## Code availability

The versions of the tools used and their parameters are described as follows.

Flye v2.3.6-release:

flye --pacbio-raw -g 2.9g

Redbean (Wtdbg2) v2.3:

wtdbg2 -x rsII -g 2.7g -L 5000

wtpoa-cns -i ctg.lay.gz

For cynomolgus macaque, rsII was replaced by sq.

SMARTdenovo (git commit 3d9c22e25bdf4caf6c08ea1acb41ee58e52f61a8): Default parameters with consensus generation.

Minimap2 v2.10-r761 and miniasm (git commit 17d5bd12290e0e8a48a5df5afaeaef4d171aa133):

minimap2 -x ava-pb | gzip -1 >m.paf.gz

miniasm -f reads.fastq m.paf.gz >tigs.gfa

For cynomolgus macaque, only reads longer than 10 kb were considered for the assembly.

Hi-C scaffolding:

Trim Galore v0.4.5: --paired --phred33 -e 0.1 -q 30

HiC-Pro v2.11.1: default parameters

Juicer v20180805: default parameters

3d-dna v20180929: -m haploid -i 5000 -r 2

Juicebox v1.3.6

Polishing:

Pbmm2 v0.12.0:

pbmm2 align assembly.referenceset.xml reads.subreadset.xml aln.alignmentset.xml --sort -j 18 -J 18 -m 5000M

Variant Caller v2.3.2:

The consensus sequence was split into 50 parts and the arrow algorithm was executed using default parameters.

Arrow polishing was iteratively executed twice for flye and SMARTdenovo assemblies, and thrice for miniasm and redbean assemblies.

Validation:

gVolante v1.2.1

## Acknowledgements

This study was supported Drug Discovery & Development program in the Japan Agency for Medical Research and Development (AMED) under grant number JP17kk0305008, and research grants to RIKEN Preventive Medicine and Diagnosis Innovation Program and RIKEN Center for Integrative Medical Sciences from The Ministry of Education, Culture, Sports, Science and Technology (MEXT). In addition, Y.S. was supported by JSPS KAKENHI Grant Numbers 18H04127, and a Grant-in-Aid for Scientific Research on Innovative Areas “Frontier Research on Chemical Communications” [no. 17H06410] from the Ministry of Education, Culture, Sports, Science and Technology of Japan. Non-Clinical Evaluation Expert Committee in Drug Evaluation Committee in Japan Pharmaceutical Manufacturers Association (JPMA) gave us constructive comments and warm encouragements. The authors have had a lot of supports and encouragements of discussion with Dr. Takao Inoue in the National Institute of Health Sciences.Special thanks to Dr. Yoshihide Hayashizaki in RIKEN Preventive Medicine and Diagnosis Innovation Program and Dr.Yasushi Okazaki in RIKEN Center for Integrative Medical Sciences.

## Author information

### Contributions

Y.Sa., J.K., H.K., Y.M. conceived, designed and planned the project; Y.Y. managed the project; V.J., Y.Sa. assembled and analyzed the genome; K.M. constructed the Hi-C library; O.N., V.J., performed Hi-C scaffolding; O.N., K.M., V.J., S.K. analyzed Hi-C scaffolding results; V.J., O.N. resolved misjoins in the Hi-C scaffolds; V.J., O.N., K.M., S.K., N.K., H.K. analyzed the data and performed technical validation; M.N., T.T., Y.Se., E.S. dissected tissues for genomic DNA; S.N., M.E., E.S. contributed to the health qualification of the monkeys; V.J. handled data submission; V.J., O.N., M.K., S.K., H.K., Y.S. wrote the manuscript; All authors have read and approved the manuscript;

