## supplementary figure S1, supplementary figure S2, supplementary figure S3, supplementary table S1, supplementary figure S2 for "Chromosomal-scale *De novo* Genome Assemblies of Cynomolgus Macaque and Common Marmoset"

### SUMMARY

The manuscript describes the construction of chromosome-scale genome assemblies of two non-human primates, cynomolgus macaque and common marmoset. This document contains supplementary figures S1,S2, S3 and supplementary tables S1, and S2.

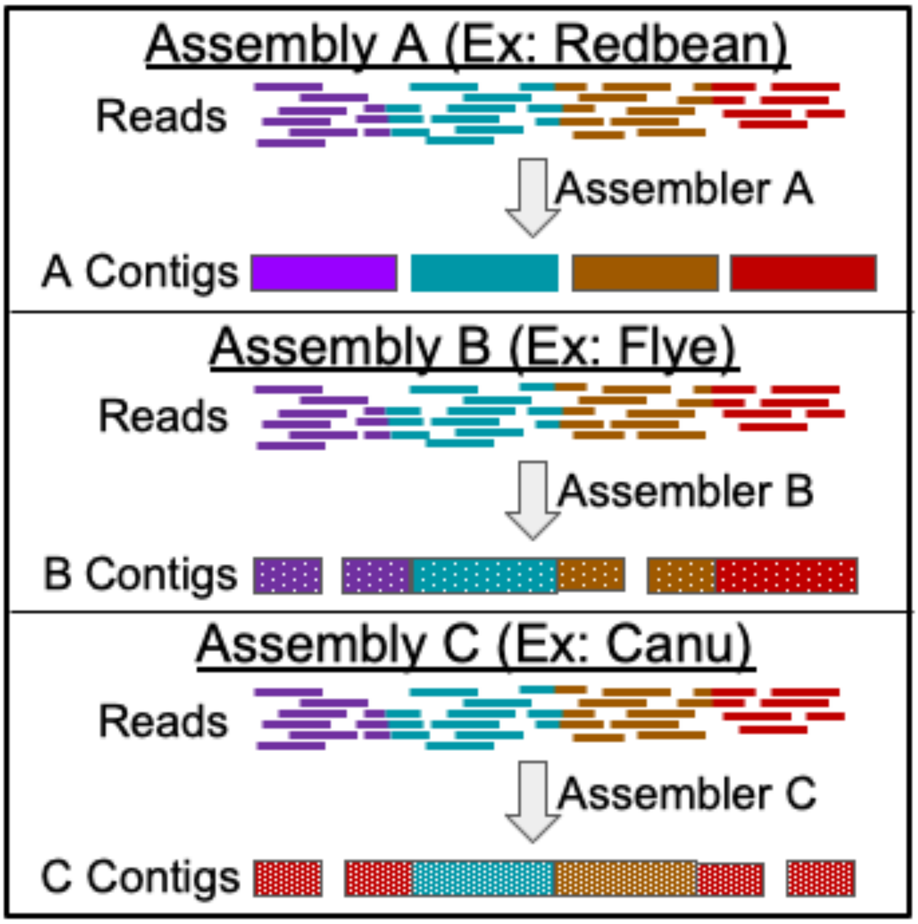


**Supplementary figure S1. *De novo* assembly from multiple assemblers**
Multiple *de novo* assembly tools employed to obtain different contiguity profiles.

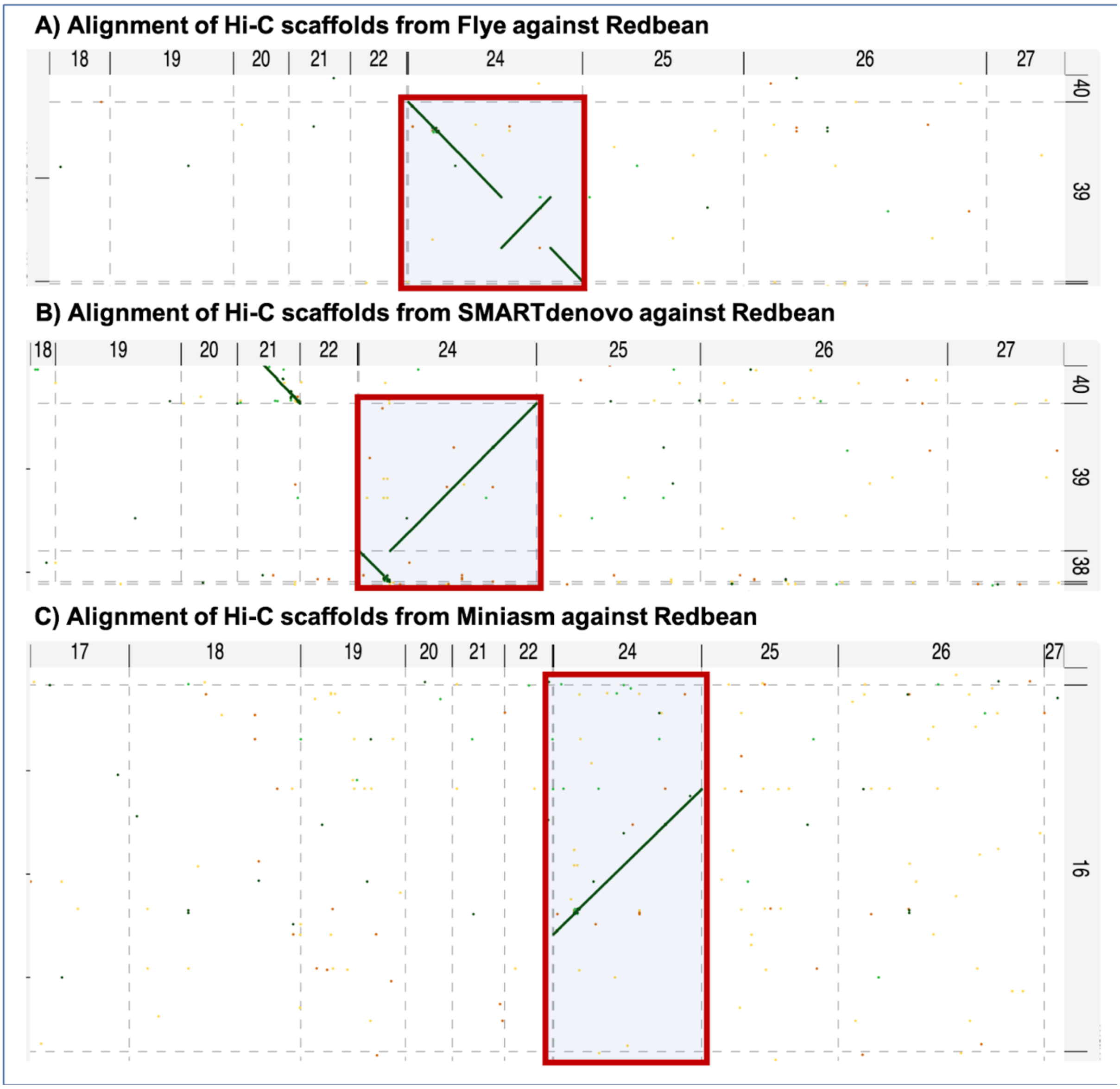


**Supplementary figure S2. Misjoins in Hi-C scaffolding**

Hi-C scaffolding of genome assemblies produced from different assembly tools revealed misjoins.

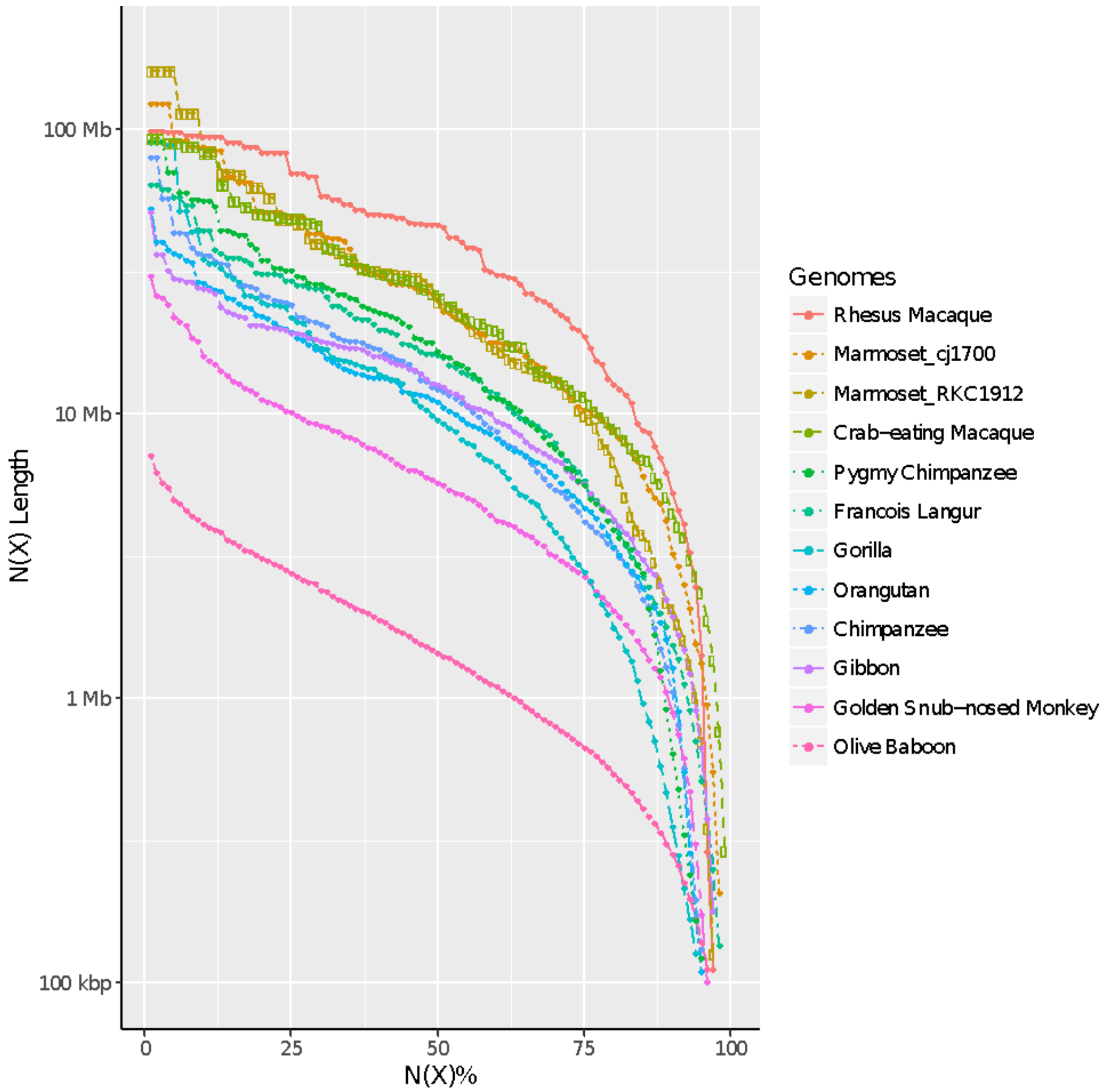


**Supplementary figure S3. Contiguity plots of non-primate genome assemblies**
N(X) plots comparing the contiguity profiles of publicly available non-human primate genome assemblies.

**Supplementary table S1: Proportion of the Hi-C libraries**

| Read pair category | Common marmoset | Cynomolgus macaque |
|---|---|---|
| Unique paired alignments | 75.2% | 67.1% |
| Valid Hi-C pairs | 72.9% | 65.4% |
| Dangling end pairs | 1.0% | 0.9% |
| Re-ligation pairs | 1.2% | 0.8% |
| Self circle pairs | 0.1% | 0.1% |
| Single-end pairs | 0.0% | 0.0% |
| Filtered pairs | 0.0% | 0.0% |
| Dumped pairs | 0.0% | 0.0% |
| Unmapped pairs | 0.6% | 1.2% |
| Low quality pairs | 0.0% | 0.0% |
| Multiple pairs alignments | 17.4% | 21.0% |
| Pairs with singleton | 6.7% | 10.6% |
| Low quality singleton | 0.0% | 0.0% |
| Unique singleton alignments | 0.0% | 0.0% |
| Multiple singleton alignments | 0.0% | 0.0% |

**Supplementary table S2. Gaps in cj1700 genome assembly filled by calJacRKC1912**

| FASTA ID of cj1700 | Total gaps (Ns) | Gaps filled by calJacRKC1912 | % of gaps filled |
|---|---|---|---|
| NC_048383.1 | 34 | 13 | 38.24 |
| NC_048384.1 | 25 | 22 | 88.00 |
| NC_048385.1 | 8 | 5 | 62.50 |
| NC_048386.1 | 20 | 5 | 25.00 |
| NC_048387.1 | 21 | 7 | 33.33 |
| NC_048388.1 | 13 | 9 | 69.23 |
| NC_048389.1 | 11 | 7 | 63.64 |
| NC_048390.1 | 4 | 3 | 75.00 |
| NC_048391.1 | 14 | 7 | 50.00 |
| NC_048392.1 | 14 | 7 | 50.00 |
| NC_048393.1 | 11 | 10 | 90.91 |
| NC_048394.1 | 16 | 4 | 25.00 |
| NC_048395.1 | 18 | 13 | 72.22 |
| NC_048396.1 | 18 | 8 | 44.44 |
| NC_048397.1 | 11 | 4 | 36.36 |
| NC_048398.1 | 8 | 1 | 12.50 |
| NC_048399.1 | 6 | 4 | 66.67 |
| NC_048400.1 | 12 | 4 | 33.33 |
| NC_048401.1 | 7 | 2 | 28.57 |
| NC_048402.1 | 6 | 5 | 83.33 |
| NC_048403.1 | 11 | 8 | 72.73 |
| NC_048404.1 | 26 | 17 | 65.38 |
| NC_048405.1 | 32 | 9 | 28.13 |
| All pseudo-chromosomes | 346 | 174 | 50.29 |
